# Automated methods enable direct computation on phenotypic descriptions for novel candidate gene prediction

**DOI:** 10.1101/689976

**Authors:** Ian R. Braun, Carolyn J. Lawrence-Dill

## Abstract

Natural language descriptions of plant phenotypes are a rich source of information for genetics and genomics research. We computationally translated descriptions of plant phenotypes into structured representations that can be analyzed to identify biologically meaningful associations. These representations include the EQ (Entity-Quality) formalism, which uses terms from biological ontologies to represent phenotypes in a standardized, semantically-rich format, as well as numerical vector representations generated using Natural Language Processing (NLP) methods (such as the bag-of-words approach and document embedding). We compared resulting phenotype similarity measures to those derived from manually curated data to determine the performance of each method. Computationally derived EQ and vector representations were comparably successful in recapitulating biological truth to representations created through manual EQ statement curation. Moreover, NLP methods for generating vector representations of phenotypes are scalable to large quantities of text because they require no human input. These results indicate that it is now possible to computationally and automatically produce and populate large-scale information resources that enable researchers to query phenotypic descriptions directly.

## 2 Background

Phenotypes encompass a wealth of important and useful information about plants, potentially including states related to fitness, disease, and agricultural value. They comprise the material on which natural and artificial selection act to increase fitness or to achieve desired traits, respectively. Determining which genes are associated with traits of interest and understanding the nature of these relationships is crucial for manipulating phenotypes. When causal alleles for phenotypes of interest are identified, they can be selected for in populations, targeted for deletion, or employed as transgenes to introduce desirable traits within and across species. The process of identifying candidate genes and specific alleles associated with a trait of interest is called candidate gene prediction.

Genes with similar sequences often share biological functions and therefore can create similar phenotypes. This is one reason sequence similarity search algorithms like BLAST are so useful for candidate gene prediction (Altschul et al., 1990). However, similar phenotypes can also be attributed to the function of genes that have no sequence similarity. This is how protein-coding genes that are involved in different steps of the same metabolic pathway or transcription factors involved in regulating gene expression contribute to shared phenotypes. For example, knocking out any one of the many genes involved in the maize anthocyanin pathway can result in pigment changes (reviewed in Sharma et al., 2011). This concept is modelled in Figure 1, where, notably, the sequence-based search with Gene 1 as a query can only return genes with similar sequences, but querying for similar phenotypes to those associated with Gene 1 returns many additional candidate genes.

**Figure 1.**
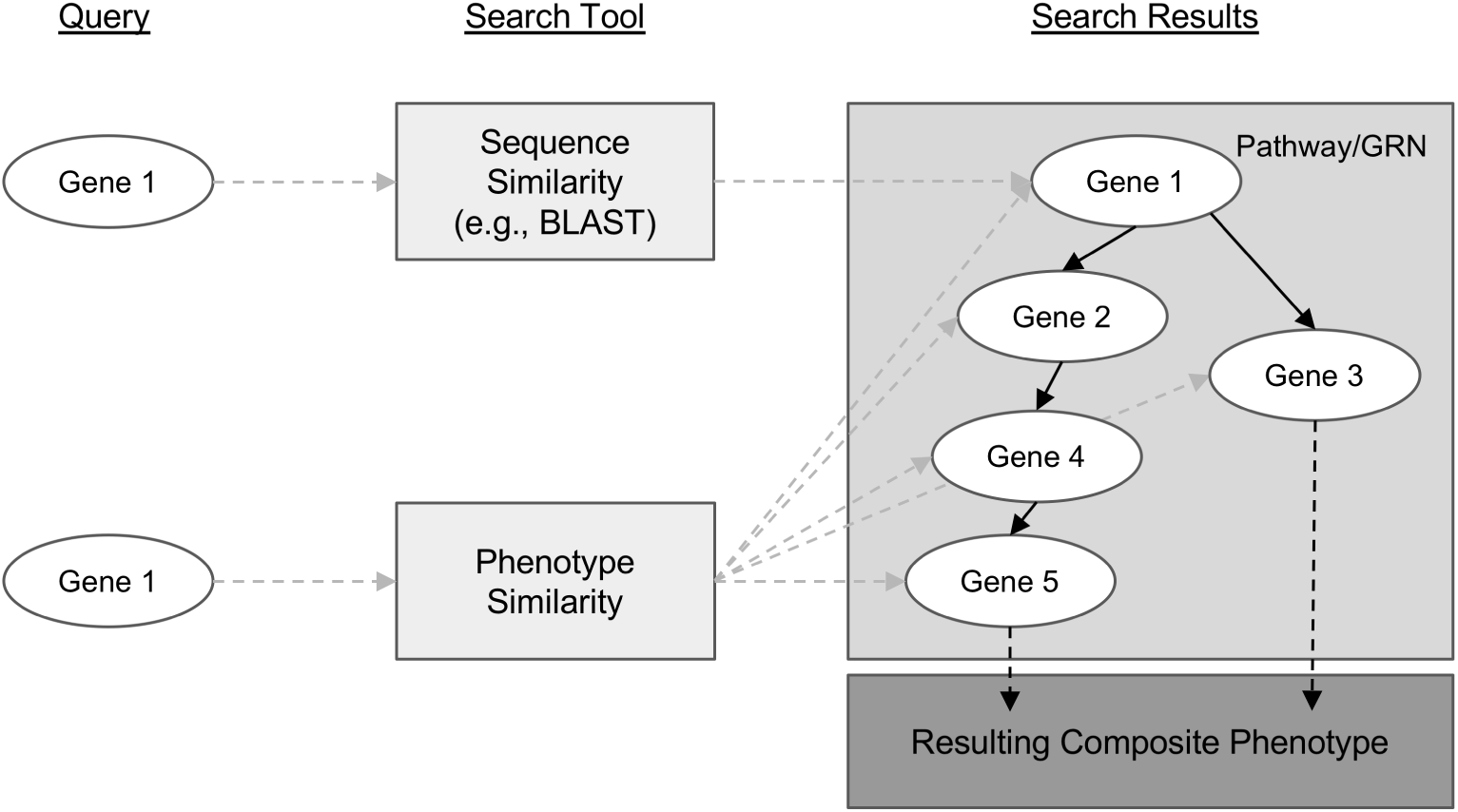
Conceptual comparison of querying with a gene sequence or its associated phenotypic description. Genes are shown as white ovals. Methods of searching for related genes are shown as light gray boxes. Gray dashed arrows indicate the path from the query gene to the set of genes that are returned from the search. Solid black arrows indicate relationships between genes in a biological pathway or gene regulatory network. Dashed black arrows indicate relationships between the pathway or network and the resulting phenotype.

High-throughput and computational phenotyping methods are largely sensor and image-based (Fahlgren et al., 2015). These methods can produce standardized datasets such that, for example, an image can be analyzed, data can be extracted, and those data can be interrogated (Gehan et al., 2017; Green et al., 2012; Miller et al., 2017). However, while such methods are adept at comparing phenotypic information between plants that are physically similar, they are limited in their ability to transfer this knowledge between physically dissimilar species. For example, traits such as leaf angle vary greatly among different species, and therefore cannot be compared directly. Moreover, where shared pathways and processes are conserved across broad evolutionary distances, it can be hard to identify equivalent phenotypes. McGary et al. (2010) call these non-obvious shared phenotypes phenologs. Between species, phenologs may present as equivalent properties in disparate biological structures (Braun et al., 2018). For example, Arabidopsis KIN-13A mutants and mouse KIF2A mutants both show increased branching in single-celled structures, but with respect to neurons in mouse (Homma et al., 2003) and with respect to trichomes in Arabidopsis (Lu et al., 2004). Taken together, the ability to compute on phenotypic descriptions to identify phenologs within and across species has the potential to aid in the identification of novel candidate genes that cannot be identified by sequence-based methods alone and that cannot be identified via image analysis.

In order to identify phenologs, some methods rely on searching for shared orthologs between causal gene sets (McGary et al., 2010; Woods et al., 2013). For example, McGary et al. (2010) identified a phenolog relationship between ‘abnormal heart development’ in mouse and ‘defective response to red light’ in Arabidopsis by identifying four orthologous genes between the sets of known causal genes in each species. However, these methods are not applicable when the known causal gene set for one phenotype or the other is small or non-existent. In these cases, using natural language descriptions to identify phenologs avoids this problem by relying only on characteristics of the phenotypes, per se. These phenotypic descriptions are a rich source of information that, if leveraged to identify phenolog pairs, can enable identification of novel candidate genes potentially involved in generating phenotypes beyond what has already been described.

Unfortunately, computing on phenotype descriptions is not straightforward. Text descriptions of phenotypes present in the literature and in online databases are irregular because natural language representations of even very similar phenotypes can be highly variable. This makes reliable quantification of phenotype similarity particularly challenging (Thessen et al., 2012; Braun et al., 2018). To represent phenotypes in a computable manner, researchers have recently begun to translate and standardize phenotype descriptions into Entity-Quality (EQ) statements composed of ontology terms, where an Entity (e.g., ‘leaf’) is modified by a Quality (e.g., ‘increased length’; Mungall et al., 2010)^1^. Using this formalism, complex phenotypes are represented by multiple EQ statements. For example, multiple EQ statements are required to represent dwarfism, where the entity and quality pairs (‘plant height’, ‘reduced’) and (‘leaf width’, ‘increased’) may be used, among others. Each of these phenotypic components of the more general phenotype is termed a ‘phene’. Because both entities and qualities are represented by terms from biological ontologies (fixed vocabularies arranged as hierarchical concepts in a directed acyclic graph), quantifying the similarity between two phenotypes that have been translated to EQ statements can be accomplished using graph-based similarity metrics (Hoehndorf et al., 2011; Slimani, 2013). Such techniques for estimating semantic similarity based on arranging concepts hierarchically in a graph have long been employed in the field of natural language processing (NLP; e.g., Resnik, 1999).

Oellrich, Walls et al. (2015) developed Plant PhenomeNET, an EQ statement-based resource primarily consisting of a phenotype similarity network containing phenotypes across six different model plant species, namely Arabidopsis (*A. thaliana*), maize (*Z. mays* ssp. *mays*), tomato (*S. lycopersicum*), rice (*O. sativa*), Medicago (*M. truncatula*), and soybean (*G. max*). Their analysis demonstrated that the method developed by Hoehndorf et al. (2011) could be used to recover known genotype to phenotype associations for plants. The authors found that highly similar phenotypes in the network (phenologs) were likely to share causal genes that were orthologous or involved in the same biological pathways. In constructing the network, text statements comprising each phenotype were converted by hand into EQ statements primarily composed of terms from the Phenotype and Trait Ontology (PATO; Gkoutos et al., 2005), Plant Ontology (PO; Cooper et al., 2013), Gene Ontology (GO; Ashburner et al., 2000), and Chemical Entities of Biological Interest (ChEBI; Hastings et al., 2013) ontology.

The success of this plant phenotype pilot project was encouraging, but to scale up to computing on all available phenotypic data for each of the six species was not a reasonable goal given that curating data for this pilot project took a good deal of time and covered only phenotypes of dominant alleles for 2,747 genes across the six species. More specifically, human translation of text statements into EQ statements is the most time consuming aspect of generating phenotype similarity networks using this method. Automation of this translation promises to increase the rate at which such networks can be generated and expanded. Notable efforts to automate this process include Semantic Charaparser (Cui, 2012, Cui et al., 2015), which extracts characters (entities) and their corresponding states (qualities) after a curation step that involves assigning terms to categories, and then mapping these characters and states to EQ statements constructed from input ontologies. Other existing annotation tools such as NCBO Annotator (Musen et al., 2012) and NOBLE Coder (Tseytlin et al., 2016) are fully automated, relying only on input ontologies. Both map words in the input text to ontology terms without imposing an EQ statement structure. State-of-the-art machine learning approaches to annotating text with ontology terms also have been developed (Hailu et al., 2019). These can be trained using a dataset such as the Colorado Richly Annotated Full-Text corpus (CRAFT; Bada et al., 2012), but are not readily transferable to ontologies that are not represented in the training set.

In addition to using ontology-based methods, similarity between text descriptions of phenotypes can also be quantified using NLP techniques such as treating each description as a bag-of-words and comparing the presence or absence of those words between descriptions, or using neural network-based tools such as Doc2Vec to embed descriptions into abstract high-dimensional numerical vectors between which similarity metrics can then be easily applied (Le and Mikolov, 2014). Conceptually, this process involves converting natural language descriptions into locations in space, such that descriptions that are near each other are interpreted as having high similarity and those that are distant have low similarity.

In this work, we demonstrate that automated techniques for generating computable representations of natural language can be applied to a dataset of phenotypic descriptions in order to generate biologically meaningful phenotype similarity networks. See Figure 2 for an overview of how phenotype similarity networks are computationally generated as an output when text descriptions are provided as the input. We first show that these computational techniques are limited in their capability to exactly reproduce the annotations and corresponding phenotype similarity networks generated with hand-curation. However, we subsequently show that the hand-curated network does not outperform networks built with purely computational approaches on dataset-wide tasks of biological relevance, such as organizing genes by function and predicting membership in biochemical pathways. Most importantly, we discuss how we can now use these computational approaches to automatically generate new datasets necessary to identify phenotypic similarities and predict gene function within and across species without requiring the use of time-consuming and costly hand-curation.

**Figure 2.**
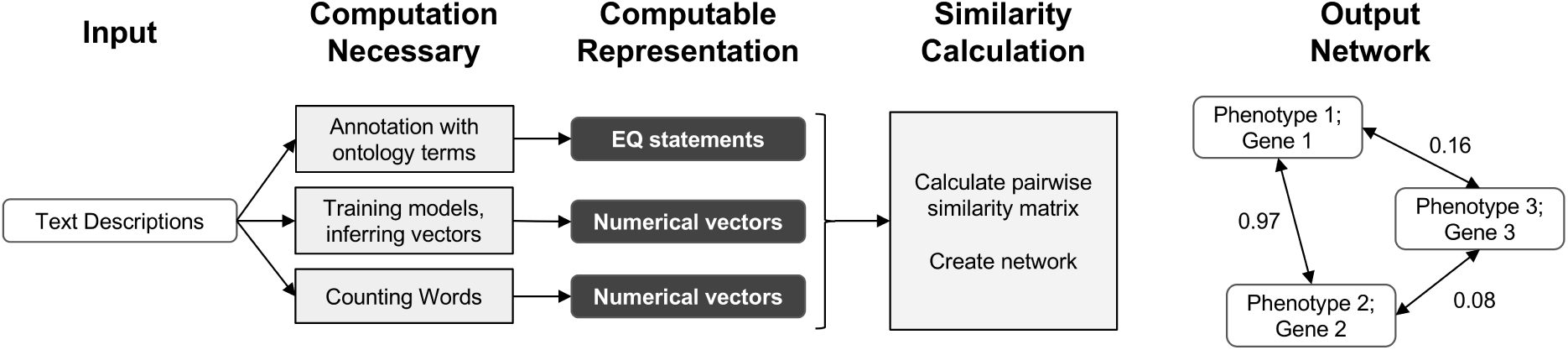
Overview of computational pipelines used here to generate phenotype similarity networks from text descriptions of phenotypes. Rounded white rectangles represent data in the form of text descriptions as input, or network nodes as output. Rounded black rectangles represent the intermediate data forms that are computable representations of text descriptions. These allow for quantitative similarity metrics to be applied. Gray rectangles represent computational methods carried out at each step. Single-headed arrows represent flow of data through each pipeline. Double-headed arrows represent edges between nodes in resulting similarity networks. Values next to double-headed arrows indicate magnitude of phenotype similarity. One output network is created for each computable representation, but only one example is shown here.

## 3 Methods

### 3.1 Dataset of Phenotypic Descriptions and Curated EQ Statements

The pairwise phenotype similarity network described in Oellrich, Walls et al. (2015) was built based on a dataset of phenotype descriptions across six different model plant species (*A. thaliana, Z. mays* ssp. *mays, S. lycopersicum, O. sativa, M. truncatula*, and *G. max*). In that work, each phenotype description was split into one or more atomized statements describing individual phenes, each of which mapped to exactly one curated EQ statement (Table 1). The EQ statements in this dataset were primarily built from terms present in PATO, PO, GO, and ChEBI. For this work, we used this existing dataset as the source of genes and associated phenotypic descriptions on which to test automated methods for assessing similarity networks between phenotypes and using the resulting phenotype similarity networks to perform comparative analyses across the whole dataset to predict gene function.

**Table 1.** Description of the Oellrich, Walls et al. (2015) dataset in terms of number of phenotype descriptions, phene descriptions, and EQ statements.

| Species | Phenotypes | Phenes<sup>1</sup> | EQ statements<sup>2</sup> |
| --- | --- | --- | --- |
| Arabidopsis | 1385 | 5172 | 5172 |
| Maize | 117 | 373 | 373 |
| Tomato | 90 | 269 | 269 |
| Rice | 86 | 340 | 340 |
| Medicago | 40 | 149 | 149 |
| Soybean | 24 | 61 | 61 |
| Example gene:<br>Arabidopsis<br>*PKS2*<br>(AT1G14280.1) | Phenotype: short<br>hypocotyl and<br>expanded cotyledon<br>under hourly far red<br>pulses | Phene 1: short<br>hypocotyl<br>Phene 2: expanded<br>cotyledon | PO:0020100 (hypocotyl) +<br>PATO:0000574 (decreased<br>length)<br>PO:0020030 (cotyledon) +<br>PATO:0000586 (increased<br>size) |

<sup>1</sup> Also referred to as ‘atomized statements’.
<sup>2</sup> Each EQ statement represents a single specific phene.

### 3.2 Computationally Generating EQ Statements from Phenotypic Descriptions

For each phenotype and phene description in the dataset, we computationally generated corresponding EQ statements without human interaction. To accomplish this, terms were first annotated to each text description, and then combined to form complete EQ statements. Two different existing computational tools and a simple machine learning technique were used to map ontology terms to text descriptions. Specifically, these were NCBO Annotator and NOBLE Coder, which are tools for matching ontology terms to specific words in text, and a Naïve Bayes bag-of-words classifier, which assign terms to descriptions based on the observed frequencies of term-word co-occurrence in a training data set. The Oellrich, Walls et al. (2015) dataset of descriptions and curated EQ statements was split into four groups such that any three groups of the dataset were used to a train a Naïve Bayes model that was then applied to the remaining group. The result of applying these three annotations methods was a set of ontology terms from PATO, PO, GO, and ChEBI assigned to each text description. Terms were then combined to form full EQ statements by assigning default root terms where none were matched, such as the entity term *whole plant* (PO:0000003), and organizing the matched terms into the different roles of the EQ statement by removing overlapping terms and automatically applying compositional rules used by curators in Oellrich, Walls et al. (2015). As an example, these rules include the fact that ChEBI terms cannot be the primary entity. The EQ statements were scored based on how well the terms aligned with the text description they were annotated to, so that the closest-matching EQ statements for each text description were output and used downstream to generate phenotype similarity networks. See the Supplemental Methods section for a more detailed description of this process.

### 3.3 Computationally Generating Numerical Vectors from Phenotypic Descriptions

In addition to generating EQ statements for each phenotype and phene description in the dataset, Doc2Vec was used for generating numerical vectors for each description. A model pre-trained on Wikipedia was used (Lau and Baldwin, 2016). In these document embeddings, positions within the vector do not refer to the presence of specific words but rather abstract features learned by the model. A size of 300 was used for each vector representation, which is the fixed vector size of the pre-trained model. In addition, vectors were generated for each description using bag-of-words and set-of-words representations of the text. For these methods, each position within the vector refers to a particular word in the vocabulary. Each vector element with bag-of-words refers to the count of that word in the description, and each vector element with set-of-words is a binary value indicating presence or absence of the word. In cases where phene descriptions were used instead of phenotype descriptions, the descriptions were concatenated prior to embedding to obtain a single vector.

### 3.4 Creating Gene and Phenotype Networks

Oellrich, Walls et al. (2015) developed a network with phenotypes as nodes and similarity between them as edges for all the phenotypes in the dataset. For each type of text representations that we generated with computational methods, comparable networks were constructed. For EQ statement representations, Jaccard similarity either taking the structure and order of terms in the EQ statement into account (referred to as metric S_1_) or ignoring the structure and treating the ontology terms in the EQ statement as an unordered set (referred to as metric S_2_) were used to determine edge values. See the Supplemental Methods section for a more detailed description of these similarity metrics. For vector representations generated using Doc2Vec and bag-of-words, cosine similarity was used. For the vector representations generating using set-of-words, Jaccard similarity was used. These networks are considered to be simultaneously gene and phenotype similarity networks, because each phenotype in the dataset corresponds to a specific causal gene, and a node in the network represents both that causal gene and its cognate phenotype. However, two phenotype descriptions corresponding to the same gene are retained as two separate nodes in the network, so while each node represents a unique gene/phenotype pair, a single gene may be represented within more than one node.

## 4 Results

### 4.1 Performance of Computational Methods in Reproducing Hand-Curated Annotations

We tested the ability of computational semantic annotation methods to assign ontology terms similar to those selected by curators to phenotype and phene descriptions in the Oellrich, Walls et al. (2015) dataset. Specifically, the ontology terms mapped by each method to a particular description were compared against the terms present in the EQ statement(s) that were created by hand-curation for that same description. Metrics of partial precision (*PP*) and partial recall (*PR*), as well as the harmonic mean of these values (*PF*_1_) as a summary statistic were used to evaluate performance (Table 2). Metrics *PP* and *PR* were applied as in Dahdul et al., (2018); see the Supplemental Methods section for a detailed description of these metrics.

**Table 2.** Performance metrics for semantic annotation methods.

| Annotator | Ontology | $n^1$ | Phenotype Description | | | Phene Descriptions | | |
| --- | --- | --- | --- | --- | --- | --- | --- | --- |
| | | | $PP^3$ | $PR^3$ | $PF_1^3$ | $PP^3$ | $PR^3$ | $PF_1^3$ |
| NOBLE Coder (precise) | PATO | 7882 | 0.641 | 0.627 | 0.634 | 0.601 | 0.572 | 0.586 |
|  | PO | 5634 | 0.622 | 0.380 | 0.472 | 0.546 | 0.294 | 0.382 |
|  | GO | 1505 | 0.514 | 0.521 | 0.517 | 0.510 | 0.514 | 0.512 |
| NOBLE Coder (partial) | PATO | 7882 | 0.412 | 0.748 | 0.532 | 0.375 | 0.689 | 0.486 |
|  | PO | 5634 | 0.309 | 0.758 | 0.439 | 0.269 | 0.659 | 0.382 |
|  | GO | 1505 | 0.102 | 0.846 | 0.182 | 0.091 | 0.839 | 0.165 |
| NCBO Annotator | PATO | 7882 | 0.640 | 0.619 | 0.629 | 0.598 | 0.563 | 0.580 |
|  | PO | 5634 | 0.550 | 0.259 | 0.352 | 0.458 | 0.170 | 0.248 |
|  | GO | 1505 | 0.478 | 0.433 | 0.454 | 0.480 | 0.424 | 0.450 |
|  | ChEBI | 775 | 0.429 | 0.888 | 0.579 | 0.431 | 0.913 | 0.586 |
| Naïve Bayes Classifier | PATO | 7882 | 0.517 | 0.394 | 0.447 | 0.642 | 0.484 | 0.552 |
|  | PO | 5634 | 0.474 | 0.258 | 0.334 | 0.636 | 0.429 | 0.512 |
|  | GO | 1505 | 0.091 | 0.073 | 0.081 | 0.155 | 0.157 | 0.156 |
|  | ChEBI | 775 | 0.035 | 0.031 | 0.033 | 0.001 | 0.001 | 0.001 |
| Aggregate Annotations<sup>2</sup> | PATO | 7882 | 0.412 | 0.798 | 0.543 | 0.383 | 0.815 | 0.522 |
|  | PO | 5634 | 0.351 | 0.809 | 0.489 | 0.304 | 0.831 | 0.445 |
|  | GO | 1505 | 0.107 | 0.839 | 0.190 | 0.090 | 0.839 | 0.163 |
|  | ChEBI | 775 | 0.366 | 0.890 | 0.519 | 0.305 | 0.913 | 0.457 |

<sup>1</sup> The number of terms from a given ontology in all curated EQ statements in the dataset.
<sup>2</sup> Annotation set formed by taking the union of the annotations from all other methods.
<sup>3</sup> Metrics are partial precision and recall ( $PP$ , $PR$ ; Dahdul et al., 2018) and their harmonic mean ( $PF_1$ ).

NOBLE Coder and NCBO Annotator generally produced semantic annotations more similar to the hand-curated dataset using phenotype descriptions as input than using the set of phene descriptions as input, a result consistent across ontologies. We considered this to be counter-intuitive, because the phene descriptions are more directly related to the individual EQ statements in terms of semantic content. However, the set of target ontology terms considered correct is larger in the case of the phenotype descriptions because this set of terms includes all terms in any EQ statements derived from that phenotype rather than a single EQ statement, which could contribute to this measured increase in both partial recall and partial precision. Accounting for synonyms and related words generated through Word2Vec models increased *PR* in the case of specific annotation methods as the threshold for word similarity was decreased (from 1.0 to 0.5) but did not increase *PF*_1_ in any instance due to the corresponding losses in *PP* (Supplemental Figure 1).

NOBLE Coder and NCBO Annotator performed comparably in the case of each type of text description and ontology, with NOBLE Coder using the precise matching parameter slightly out-performing the other annotation method with respect to these particular metrics for these particular descriptions. Both outperformed the Naïve Bayes classifier, for which performance dropped significantly for the ontologies with smaller relative representation in the dataset (GO and ChEBI) as might be expected. When the results were aggregated, the increase in partial recall for PATO, PO, and GO terms relative to the maximum recall achieved by any individual method indicates that the curated terms that were recalled by each method were not entirely overlapping. This is as expected given that different methods used for semantic annotation recalled target (curated) ontology terms to different degrees, as measured by Jaccard similarity of a given target term to the closest predicted term annotated by that particular method. These sets of obtained similarities to target terms were comparable between NCBO Annotator and NOBLE Coder (*ρ* = 0.84 with phene descriptions and *ρ* = 0.86 with phenotype descriptions), and dissimilar between either of those methods and the Naïve Bayes classifier (*ρ <* 0.10 in both cases for either type of description), using Spearman rank correlation adjusted for ties.

These results indicate that automated annotation methods (NCBO Annotator, NOBLE Coder, Naïve Bayes classifier) do not reproduce the exact same ontology term annotations selected by hand-curation for each phenotypic description, as expected. Given this result, we next assessed how these differences between the hand-curated annotations and computationally generated annotations translated into differences between the phenotype similarity networks based on these annotations.

### 4.2 Comparing Computational Networks to the Hand-Curated Network

Oellrich, Walls et al. (2015) developed a network with phenotype/gene pairs as nodes and similarity between them as edges for all phenotypes in the dataset. In this work, comparable networks were constructed for the same dataset using a number of computational approaches for representing phenotype and phene descriptions and for predicting similarity. For the purposes of this assessment, the network built from hand-curated EQ statements and described in Oellrich, Walls et al. (2015) is considered the gold standard against which each network we produced is compared. The computational and gold standard networks were compared using the F1 metric to assess similarity in predicted phenolog pairs at a range of *k* values where *k* is the allowed number of phenolog pairs predicted by the networks (the *k* most highly valued edges). Results are reported through *k*=583,971, which is the number of non-zero similarities between phenotypes in the gold standard network, and were repeated using phenotype descriptions and phene descriptions as inputs to the computational methods (Figure 3). The simplest NLP methods for assessing similarity (set-of-words and bag-of-words) consistently recapitulated the gold standard network the best using phenotype descriptions, whereas the document embedding method using Doc2Vec outperformed these methods for values of *k ≤* 200,000 based on phene descriptions. The differences in the performance of each method are robust to 80% subsampling of the phenotypes present in the dataset.

**Figure 3.**
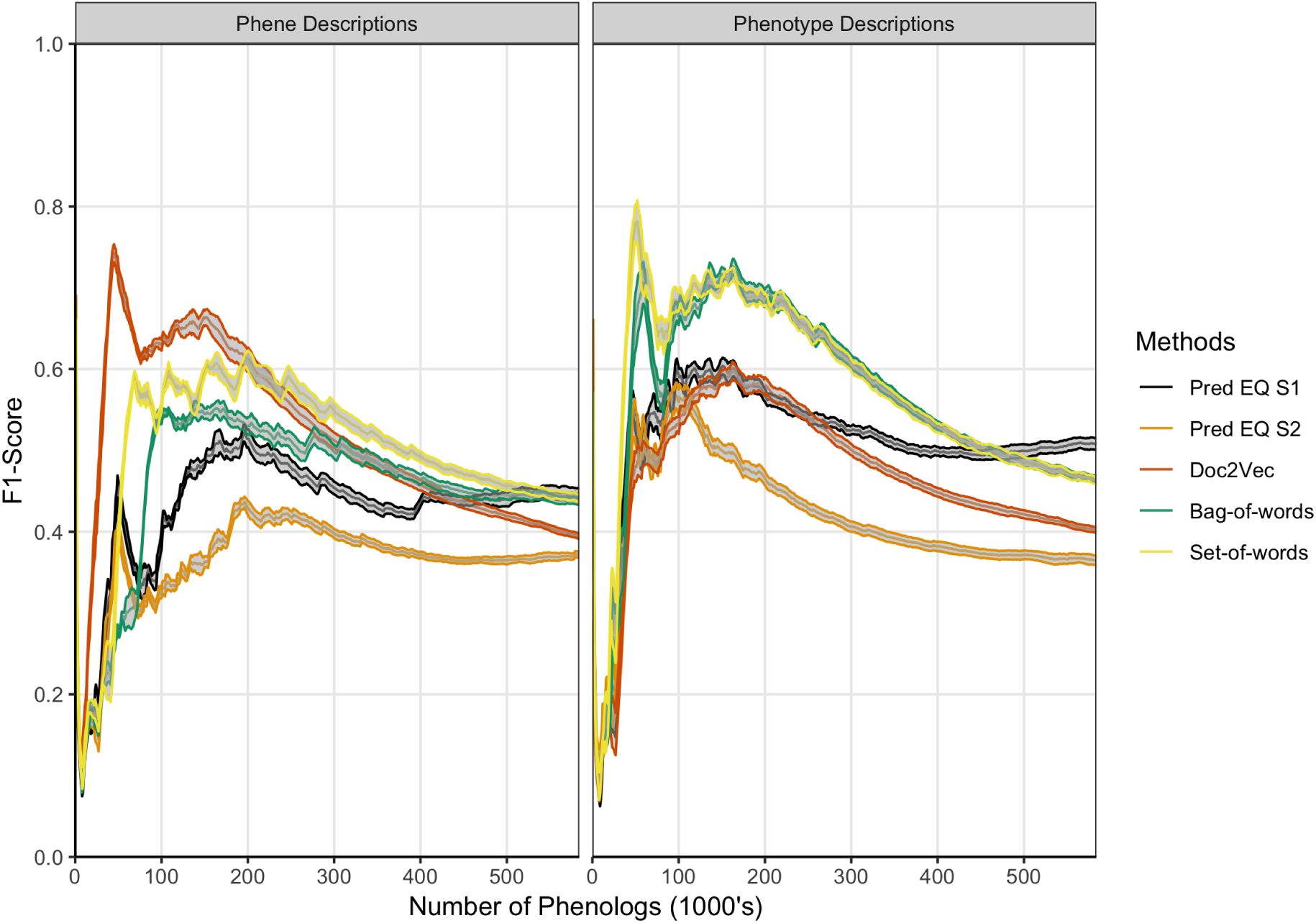
Comparison of phenolog pairs identified by predictive methods in comparison to the Oellrich, Walls et al. (2015) dataset. The *x*-axis indicates the number of phenologs pairs (highest valued edges in the phenotype similarity network) at each point. The standard deviation of resampling with 80% of the phenotypes in the dataset (network nodes) are indicated by ribbons for each method. Phene descriptions (left) or phenotype descriptions (right) were used as the text input for each particular method.

These results illustrate that computational methods do not exactly reproduce the phenotype similarity network built from the hand-curated EQ statements. However, this does not necessarily mean that the hand-curated network is inherently more biologically meaningful. To assess whether this is true, we next compared how the hand-curated network and each computational network performed on an independent biologically-relevant task– sorting genes into functional groups.

### 4.3 Computational Methods Outperform Hand-Curation for Gene Functional Categorization in Arabidopsis

Lloyd and Meinke (2012) previously organized a set of Arabidopsis genes with accompanying phenotype descriptions into a functional hierarchy of groups (*e.g.* ‘morphological’), classes (*e.g.* ‘reproductive’), and finally subsets (*e.g.* ‘floral’), in order from most general to most specific. See Supplemental Table 1 in Lloyd and Meinke (2012) for a full specification of this hierarchy to which the genes were assigned, and Supplemental Table 2 in Lloyd and Meinke (2012) for a mapping between genes and this hierarchical vocabulary. Oellrich, Walls et al. (2015) later used this set of genes and phenotypes to validate the quality of their dataset of hand-curated EQ statements by reporting the average similarity of phenotypes (translated into EQ statements) that belonged to the same functional subset. We used this same functional hierarchy categorization and a similar approach to assess the utility of computationally generated representations of phenotypes towards correctly categorizing the functions of the corresponding genes, and to compare this utility against that of the dataset of hand-curated EQ statements. For each class and subset in the hierarchy, the mean similarity between any two phenotypes related to genes within that class or subset (‘within’ mean) was quantified using each computable representation of interest, and compared to the mean similarity between a phenotype related to a gene within that class or subset and one outside of it (‘between’ mean), quantified in terms of standard deviation of the distribution of all similarity scores generated for each given method. The difference between the ‘within’ mean and ‘between’ mean (referred to here as the Consistency Index) for each functional category for each method indicates the ability of that method to generate strong similarity signal for phenotypes in this dataset that share that function (Figure 4). In the case of these data, most computational methods using either phene or phenotype descriptions as the input text were able to recapitulate the signal present in the network Oellrich, Walls et al. (2015) generated from hand-curated EQ statements, and the simplest NLP methods (bag-of-words and set-of-words) produced the most consistent signal.

**Figure 4.**
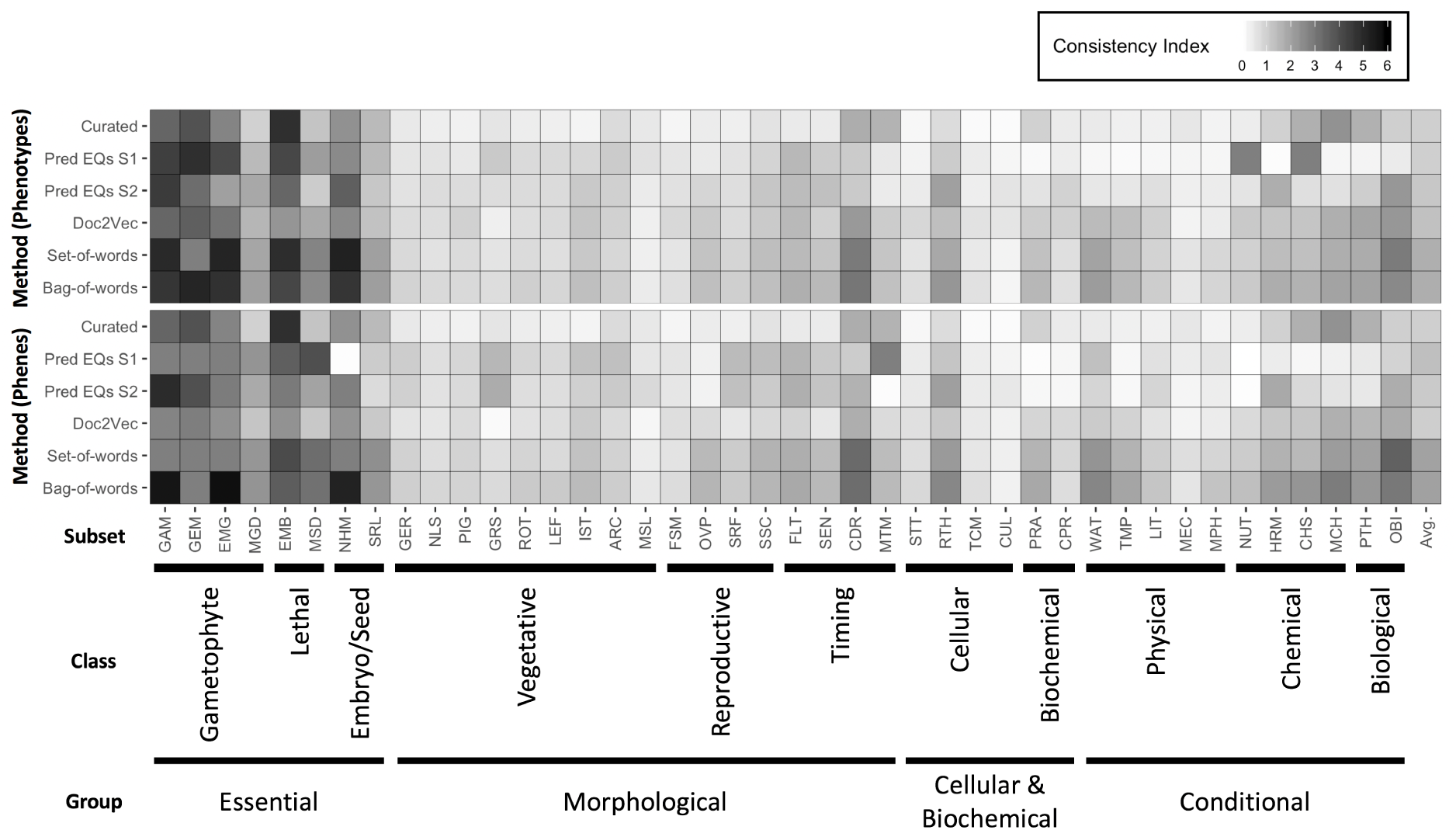
Heatmap of Consistency Index. The difference between average similarity for two phenotypes within a subset and one phenotype within and one outside, for each functional subset defined in the dataset of Arabidopsis phenotypes, and for each method of quantifying similarity between phenotypes is shown, with darker cells indicating higher consistency within a subset. Differences are measured in standard deviations of the distributions of similarities obtained for each method. The meaning of subset abbreviations are specified in Supplemental Table 1 of Lloyd and Meinke (2012). Methods are listed at left. Input text for calculating similarities between the phenotypes were either derived from phenotype descriptions (top) or phene descriptions (bottom). The far right column in the heatmap refers to an average Consistency Index for a given method across all subsets.

In order to more directly compare each method on a general classification task, networks constructed from curated EQ statements and those generated using each computational method were used to iteratively classify each Arabidopsis phenotype into classes and subsets. This was accomplished by removing one phenotype at a time and withholding the remaining phenotypes as training data, learning a threshold value from the training data, and then classifying the held out phenotype by calculating its average similarity to each training data phenotype in each class or subset, and classifying it as belonging to any category for which the average similarity to other phenotypes in that category exceeded the learned threshold. Performance on this classification task using each network was assessed using the F1 metric, where the functional category assignments for each gene reported by Lloyd and Meinke (2012) were considered to be the correct classifications (Table 3). The simplest NLP methods (bag-of-words and set-of-words) outperformed the Oellrich, Walls et al. (2015) hand-curated EQ statement network on this classification task in all cases, while using the computationally generated EQ statements or document embeddings generated with Doc2Vec only outperformed the curated EQ statement network in some cases.

**Table 3.** Evaluation (F1 scores) for each method used to categorize Arabidopsis genes by function.

| Method | Phenes |  | Phenotypes |  |
| --- | --- | --- | --- | --- |
|  | Class | Subset | Class | Subset |
| Curated EQ | 0.470 | 0.359 | 0.470 | 0.359 |
| Pred EQ S1 | 0.472 | 0.472 | 0.369 | 0.320 |
| Pred EQ S2 | 0.504 | 0.413 | 0.437 | 0.368 |
| Set-of-words | 0.613 | 0.447 | 0.587 | 0.426 |
| Bag-of-words | 0.595 | 0.423 | 0.549 | 0.409 |
| Doc2Vec | 0.455 | 0.331 | 0.486 | 0.377 |

Taken together, these results indicate that even though the computationally generated networks are significantly different than the hand-curated network (Figure 3), they generally perform equally-well *or better* on tasks related to organizing Arabidopsis genes into functional groups. We next examined how these networks compare on the task of predicting biochemical pathway membership for specific genes, both within a single species and across multiple species.

### 4.4 Computational Methods Outperform Hand-Curation for Recovering Genes Involved in Anthocyanin Biosynthesis Both Within and Between Species

Oellrich, Walls et al. (2015) illustrated the utility of using EQ statement representations of phenotypes to provide semantic information necessary to recover shared membership of causal genes in regulatory and metabolic pathways. Specifically, they showed that by querying a six-species phenotype similarity network with the *c2* (*colorless2*) gene in maize, which is involved in anthocyanin biosynthesis, genes *c1, r1*, and *b1* (*colorless1, red1, booster1*) which are also involved in anthocyanin biosynthesis in maize are recovered. Querying in this instance is defined as returning other genes in the similarity network, ranked using the maximal value of the edges connecting a phenotype corresponding to the query gene and a phenotype corresponding to each other gene in the network. There are 2,747 genes in the dataset, so querying with one gene returns a ranked list of 2,746 genes. This result was included by Oellrich, Walls et al. (2015) as a specific example of the general utility of the phenotype similarity network to return other members of a pathway or gene regulatory network when querying with a single gene. See Figure 1 for a general illustration of this concept.

To evaluate this same utility in the phenotype similarity networks we generated using computational methods and to compare their utility to that of the network from Oellrich, Walls et al. (2015) generated using hand-curated EQ statements, we first expanded the set of maize anthocyanin pathway genes to include those present in the description of the pathway given by Li et al., (2019), and listed in Supplementary Table 1 of that publication. Of those genes, 10 are present in the Oellrich, Walls et al. (2015) dataset (Table 4). Additionally, we likewise identified the set of Arabidopsis genes known to be involved in anthocyanin biosynthesis (listed in Table 1 of Appelhagen et al. (2014)) that were present in the Oellrich, Walls et al. (2010) dataset. This yielded a total of 16 Arabidopsis genes (Table 5).

**Table 4.** Maize genes involved in anthocyanin biosynthesis.

| Gene Name (Symbol) | Gene Model ID<sup>1</sup> | Category<sup>2</sup> | Encoded Protein<sup>3</sup> |
| --- | --- | --- | --- |
| colorless2 (*c2*) | **GRMZM2G422750** | Enzyme | naringenin-chalcone synthase |
| *chalcone flavonone isomerase1* (*chi1*) | GRMZM2G155329 | Enzyme | chalcone isomerase |
| *red aleurone1* (*pr1*) | GRMZM2G025832 | Enzyme | flavonoid 3'-hydroxylase (flavonoid 3'-monooxygenase) |
| *flavanone 3-hydroxylase1* (*fht1*; *F3H*) | GRMZM2G062396 | Enzyme | flavonone 3'-hydroxylase (flavonol synthase) |
| *anthocyaninless1* (*a1*) | **GRMZM2G026930** | Enzyme | dihydroflavonol 4-reductase (flavonone 4-reductase) |
| *anthocyaninless2* (*a2*) | **GRMZM2G345717** | Enzyme | anthocyanidin synthase (leucoanthocyanidin dioxygenase) |
| *bronze1* (*bz1*) | **GRMZM2G165390** | Enzyme | flavonol 3-O-glucosyltransferase |
| *bronze2* (*bz2*) | **GRMZM2G016241** | Enzyme | glutathione transferase (maleylacetoacetate isomerase) |
| *multidrug resistance associated protein3* (*mrpa3*; *ZmMrp4*) | GRMZM2G111903 | Transporter | multidrug-resistance-like-transporter |
| *scutellar node color1* (*sn1*) | GRMZM5G822829 | TF | bHLH |
| *colorless1* (*c1*) | **GRMZM2G005066** | TF | R2 R3-MYB |
| *pericarp color1 p1* | **GRMZM2G084799** | TF | R2 R3-MYB |
| *purple plant1* (*pl1*) | **GRMZM2G701063** | TF | R2 R3-MYB |
| *colored1* (*r1*) | **GRMZM5G822829** | TF | bHLH |
| *colored plant1* (*b1*) | **GRMZM2G172795** | TF | bHLH |
| *pale aleurone color1* (*pac1*) | GRMZM2G058432 | TF | WD40 |

<sup>1</sup> Gene model IDs in bold were present in the Oellrich, Walls et al. (2015) dataset.
<sup>2</sup> The abbreviation TF is short for transcription factor.
<sup>3</sup> Enzyme encoded protein names from the Plant Metabolic Network (Schlapfer et al., 2017).

**Table 5.** Arabidopsis genes involved in anthocyanin biosynthesis.

| Gene Name (Symbol) | Locus Name<sup>1</sup> | Category<sup>2</sup> | Encoded Protein<sup>3</sup> |
| --- | --- | --- | --- |
| *TRANSPARENT TESTA 4 (TT4)* | **At5g13930** | Enzyme | naringenin-chalcone synthase |
| *TRANSPARENT TESTA 5 (TT5)* | **At3g55120** | Enzyme | chalcone isomerase |
| *TRANSPARENT TESTA 6 (TT6)* | **At3g51240** | Enzyme | flavanone 3'-hydroxylase (flavonol synthase) |
| *TRANSPARENT TESTA 7 (TT7)* | **At5g07990** | Enzyme | flavonoid 3'-hydroxylase (flavonoid 3'-monooxygenase) |
| *TRANSPARENT TESTA 3 (TT3)* | **At5g42800** | Enzyme | dihydroflavonol 4-reductase (flavonone 4-reductase) |
| *TRANSPARENT TESTA 11 (TT11)*<br>*TRANSPARENT TESTA 17 (TT17)*<br>*TRANSPARENT TESTA 18 (TT18)*<br>*TANNIN-DEFICIENT SEED 4 (TDS4)* | At4g22880 | Enzyme | anthocyanidin synthase (leucoanthocyanidin dioxygenase) |
| *ARABIDOPSIS SIP1 CLADE TRIHELIX 1 (AST1)*<br>*BANYULS (BAN1)* | **At1g61720** | Enzyme | anthocyanidin reductase |
| *TRANSPARENT TESTA 14 (TT14)*<br>*TRANSPARENT TESTA 19 (TT19)* | **At5g17220** | Enzyme | glutathione transferase (maleylacetoacetate isomerase) |
| *AUTOINHIBITED H<sup>+</sup>-ATPASE (AHA10)* | At1g17260 | Enzyme | ATP-ase |
| *TRANSPARENT TESTA 10 (TT10)* | **At5g48100** | Enzyme | laccase |
| *TRANSPARENT TESTA 5 (TT15)* | **At1g43620** | Enzyme | 3β-hydroxy sterol glucosyltransferase |
| *TRANSPARENT TESTA 12 (TT12)* | **At3g59030** | Transporter | MATEefflux proton antiporter |
| *TRANSPARENT TESTA 16 (TT16)* | **At5g23260** | TF | K-box, MADS-box |
| *TRANSPARENT TESTA 1 (TT1)* | **At1g34790** | TF | C2H2 |
| *TRANSPARENT TESTA 2 (TT2)* | **At5g35550** | TF | bHLH |
| *TRANSPARENT TESTA 8 (TT8)* | **At4g09820** | TF | bHLH |
| *TRANSPARENT TESTA GLABRA 1 (TTG1)* | **At5g24520** | TF | WD40 |
| *TRANSPARENT TESTA GLABRA 2 (TTG2)* | **At2g37260** | TF | WRKY |

<sup>1</sup> Locus names in bold were present in the Oellrich, Walls et al. (2015) dataset.
<sup>2</sup> The abbreviation TF is short for transcription factor.
<sup>3</sup> Enzyme encoded protein names from the Plant Metabolic Network (Schlapfer et al., 2017).

#### 4.4.1 Recovering Anthocyanin Biosynthesis Genes Within a Single Species

Using each phenotype similarity network, each anthocyanin biosynthesis gene from one species was iteratively used as a query against the network. The rank of each other gene in the set of anthocyanin biosynthesis genes corresponding to the same species as the query was quantified. We grouped the ranks into bins of width 10 for ranks less than or equal to 50, and combined all ranks greater than 50 into a single bin. For each phenotype similarity network, the mean and standard deviation of the number of anthocyanin biosynthesis genes in each bin was calculated (Figure 5). The average number of pathway genes ranked within the top 10 across all queries was greater for all computationally generated networks than for the network built from hand-curated EQ statements, although variance across the queries was high. In general, computational networks built from predicted EQ statements performed best for this task, whereas the network built using the hand-curated EQs performed the worst. The networks constructed using the numerical vector representations (set-of-words, bag-of-words, and Doc2Vec) were intermediate in performance as a group (Figure 5).

**Figure 5.**
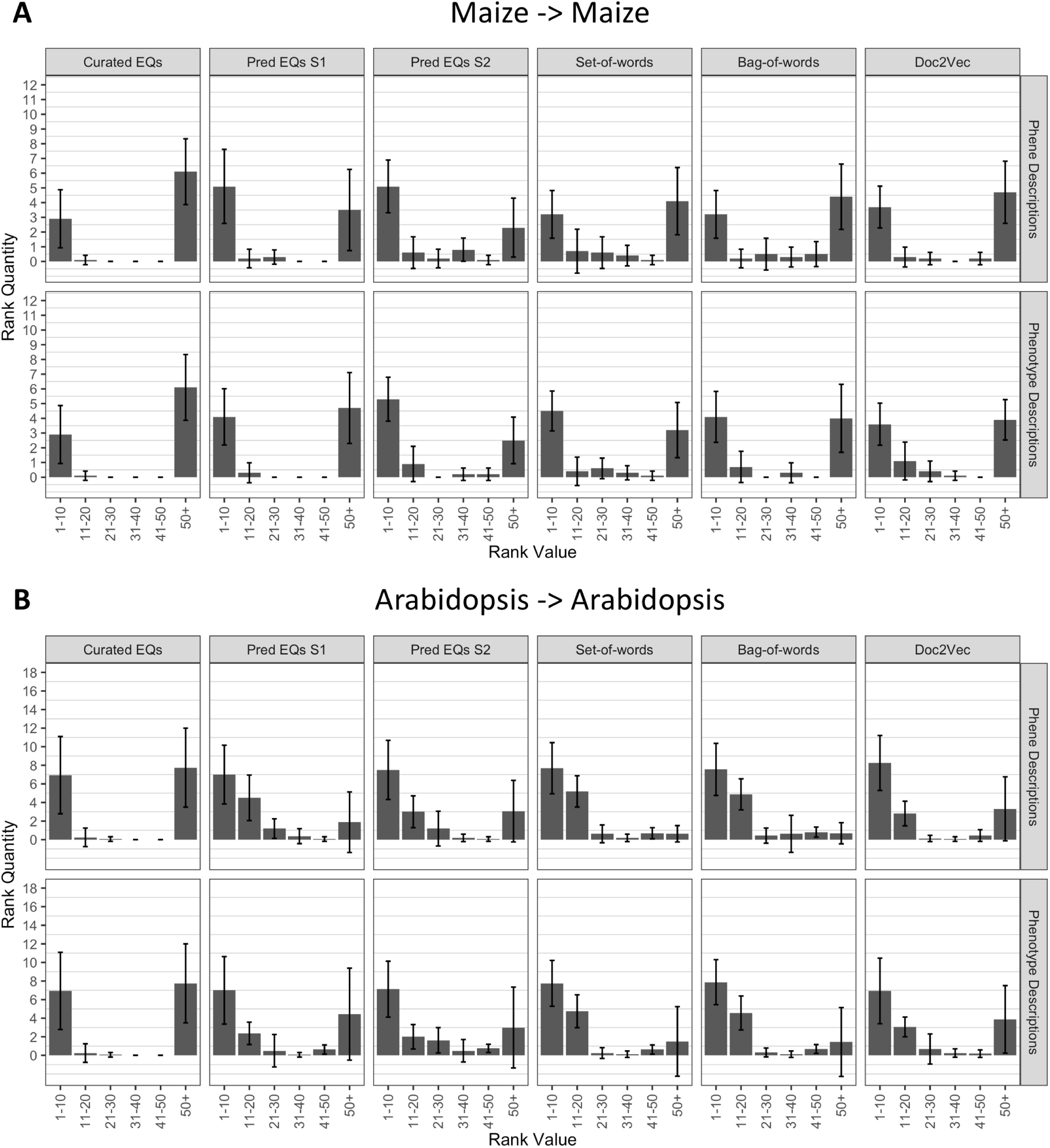
Rankings of anthocyanin biosynthesis genes in either maize (**A**) or Arabidopsis (**B**) upon querying phenotype similarity networks generated with genes from the same species. Phenotype networks are organized by the method used to generate them (columns) and by whether those methods were applied to phenotype or phene descriptions (rows). Rank value specifies a range of rankings for each bar in the plots (1-10, 11-20, etc.) and rank quantity indicates the average number of anthocyanin biosynthesis genes that were ranked in a given range over all queries. Black lines indicate plus and minus one standard deviation of the rank quantities in each range over all queries.

#### 4.4.2 Recovering Anthocyanin Biosynthesis Genes Between Two Species

To determine whether the methods performed similarly both within and across species, we repeated the analysis described in the previous section (4.4.1), but instead of quantifying the ranks of all anthocyanin biosynthesis genes from the same species as the query gene, we quantified the ranks of all anthocyanin genes that derived from the other species. In other words, Arabidopsis genes were used to query for maize genes, and maize genes were used to query for Arabidopsis genes. As shown in Figure 6, the phenotype similarity network constructed from hand-curated EQ statements did not recover (provide ranks of less than or equal to 50) any of the anthocyanin biosynthesis genes when queried with genes from the other species. Networks generated using the set-of-words and bag-of-words approaches, or with Doc2Vec, performed similarly. Networks built from computationally generated EQ statements recovered the most anthocyanin biosynthesis genes on average across the queries between species (Figure 6).

**Figure 6.**
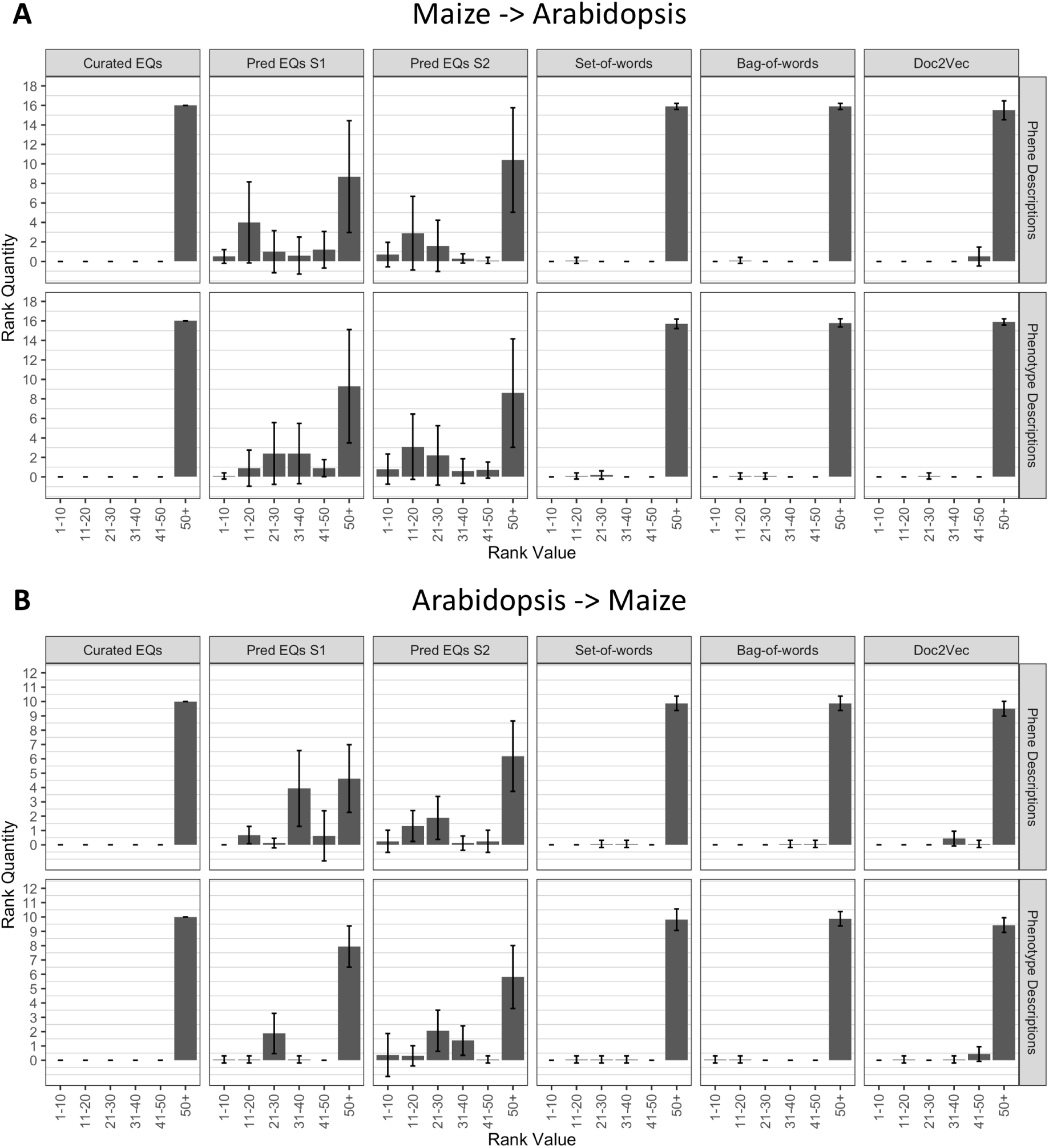
Rankings of anthocyanin biosynthesis genes in either maize (**A**) or Arabidopsis (**B**) upon querying phenotype similarity networks generated with genes from the other species. Phenotype networks are organized by the method used to generate them (columns) and by whether those methods were applied to phenotype or phene descriptions (rows). Rank value specifies a range of rankings for each bar in the plots (1-10, 11-20, etc.) and rank quantity indicates the average number of anthocyanin biosynthesis genes that were ranked in a given range over all queries. Black lines indicate plus and minus one standard deviation of the rank quantities in each range over all queries.

## 5 Discussion

### 5.1 Computationally Generated Phenotype Representations are Useful

A primary purpose for generating representations of phenotypes that are easy to compute on (EQ statements, vector embeddings, etc.) is to construct similarity networks that enable the use of one phenotype as a query to retrieve similar phenotypes. This process serves as a means of discovering relatedness between phenotypes (potential phenologs) within and across species, thus generating hypotheses about underlying genetic relatedness (reviewed in Oellrich, Walls et al., 2015).

The computational methods discussed in this work were demonstrated to only partially recapitulate the phenotype similarity network constructed by Oellrich, Walls et al. (2015) using hand-curated EQ statements (Section 4.2). Despite the limited similarity between the network built from hand-curated annotations and the computationally generated networks, the computationally generated networks performed as well or better than the hand-curated network (based on curated EQ statements) in terms of correctly organizing phenotypes and their causal genes into functional categories at multiple hierarchical levels (Section 4.3). In addition, each computationally generated network performed better than the hand-curated network for querying with either maize or Arabidopsis anthocyanin biosynthesis genes to return other anthocyanin biosynthesis genes from the same species (Section 4.4.1), a task originally used to demonstrate the utility of the phenotype similarity network constructed in Oellrich, Walls et al. (2015).

Moreover, the networks built from computationally generated EQ statements were useful for recapturing anthocyanin biosynthesis genes from a species different than the species of origin for the queried gene/phenotype pair. None of the other networks, including the network built from curated EQ statements, exhibited this utility for this task (Section 4.4.2). This particular result indicates that high accuracy of constructed EQ statements is not specifically necessary for tasks such as querying for related genes across species, because potentially inaccurate (computationally predicted) EQ statements generated a more successful network for the task. Replicating these analyses with phenotype descriptions in a different biological domain, such as vertebrates, would determine whether these results generalize to additional species groups and datasets.

Taken together, these results over this particular dataset of phenotype descriptions suggest that while the EQ statements generated through manual curation are likely the most accurate and informative computable representation of a given phenotype in specific cases, other representations generated entirely computationally with no human intervention are capable of meeting or exceeding the performance of the hand-curated annotations on dataset-wide tasks such as sorting phenotypes and genes into functional categories, as well as in the case of specific tasks such as querying with particular genes to recover other genes involved in the same pathway. Therefore, in cases where the volume of data is large, the results are understood to be predictive, and manual curation is impractical, using automated annotation methods to generate large-scale phenotype similarity networks is a worthwhile goal and can provide biologically relevant information that can be used for hypothesis generation, including novel candidate gene prediction.

### 5.2 Multiple Approaches to Representing Natural Language are Useful

EQ statement annotations comprised of ontology terms allow for interoperability with compatible annotations from varied data sources. They are also a human-readable annotation format, meaning that a knowledgeable human could fix an incorrect annotation by selecting a more appropriate ontology term (a process that is not possible using abstract vector embeddings). Their uniform structure also provides a means of explicitly querying for phenotypes involving a biological entity that is similar to some structure or process (e.g., trichomes) or matches some quality (e.g., an increase in physical size). Ontology-based annotations have the potential to increase the information attached to a phenotype (through inferring ancestral terms which are not specifically referred to in the phenotype description), but do not necessarily fully capture the detail and semantics of the natural language description.

For this reason, future representations of phenotypes in relational databases for the purpose of generating phenotype similarity networks across a large volume of phenotypes described in literature and in databases likely should include both ontology-based annotations describing the phenotypes, as well as the original natural language descriptions. Although the number of phenotypes in the dataset used here and described in Oellrich, Walls et al. (2015) is relatively small, the results of this work suggest utility of original text representations as a powerful means of calculating similarity between phenotypes, especially within a single species. Computationally generated EQ statements, which in the context of this study do not often meet the criteria for a fully logical curated EQ statement, were demonstrated to be more useful in any other approach for recovering biologically related genes across species.

Ensemble methods are often applied in the field of machine learning, where multiple methods are used to solve a problem, with a higher-level model determining which method will be most useful in solving each new instance of the problem. It is possible that such an approach could be applied to measuring similarity between phenotypes to generate a single large-scale network, where similarity values are based on the best possible method to assess the text representations of each pair of particular phenotypes.

### 5.3 Additional Challenges with EQ Statement Representation

Although ontology terms and EQ statements composed of ontology terms are an information-rich representation of phenes and phenotypes, flexibility in which terms and statements can represent a particular phenotype can limit the ability to computationally recognize true biological similarity. The graph structures of the ontologies themselves, the metrics used to assess semantic similarity, and the ambiguity inherent in both natural language and EQ statement representations of phenes and phenotypes can all potentially contribute to this problem.

As one example in the Oellrich, Walls et al. (2015) dataset used here, the phene description ‘complete loss of flower formation’ was annotated with an EQ statement whose entity is *flower development*, whereas the computationally identified entity using the methods described in this work was *flower formation*. In this instance, the Jaccard similarity between these two ontology terms was 0.286, which by comparison is less than the Jaccard similarity between *flower formation* and *leaf formation* in the context of the ontology graph. This selected example illustrates the possible discrepancies between true biological similarity and semantic similarity as measured using graph-based metrics. Although each semantic similarity metric calculates this value differently, those that use the hierarchical nature of the ontology are all constrained by the structure of the graph itself.

Variation in how humans and computational methods interpret how a phenotype as a whole should be conceptualized also has the potential to produce representations that obscure true similarity, as measured by graph-based metrics. In another example from the Oellrich, Walls et al. (2015) dataset, the phene description ‘stamens transformed to pistils’ was annotated with two different EQ statements. The first EQ statement uses the relational quality *has fewer parts of type* to indicate the absence of stamen in this phenotype, and the second uses the relational quality *has extra parts of type* to indicate the presence of pistils in this phenotype. This representation of the phenotype makes logical sense, but is not easy to generate computationally because it abstractly describes the outcome of the transformation that is explicitly present in the natural language description, and is dissimilar from computationally generated representations that focus on the explicit content (i.e., those which use the relational quality *transformed to*).

Finally, this study looked at a dataset consisting entirely of phenotypic descriptions in English, and the generalizability of these methods to other languages is not discussed. It is certainly likely that that structural differences between languages would result in differences in how certain methods of computing over descriptions in those languages perform, but such analysis is outside the scope of this work.

### 5.4 Extending this Work to the Wealth of Text Data available in Databases and the Literature

We plan to apply the methods of semantic annotation, ontology-based semantic similarity calculation, and natural language-based semantic similarity calculation to the wealth of text data available in existing plant model organism databases and biological literature. For the latter, doing so will involve the additional challenge of extracting phenotypes descriptions as well as the genes causative to those phenotypes as a separate identification and processing step. We plan to leverage existing work in the areas of named entity recognition specific to genes (Wei et al., 2015) and relation extraction, as well as existing methods for extracting information related to phenotypes such as those developed using vector-based representations of phenotype descriptions (Xing et al., 2018) and grammar-tree representations of phenotype descriptions (Collier et al., 2015). As the size of the applicable dataset is increased by these means, we will continue to analyze the performance of methods from the domains of machine learning and NLP towards constructing biologically meaningful networks from this phenotypic data, including additional techniques that were not included in the results presented here. For example, Sent2Vec (Pagliardini et al., 2018) is another technique for assessing text similarity that takes a different approach than Doc2Vec for embedding text as numerical vectors, and has been shown to perform well when trained on life science corpora (Chen et al., 2018). These next steps are anticipated to enable researchers to begin to compute on phenotype descriptions directly, and will drive a promising future for forward genetics research approaches where phenotypes can be used for novel candidate gene prediction as easily as sequence similarity searches can be used to identify putative homologs from sequence data.

## Data Availability

The dataset of phenotype and phene descriptions and the corresponding hand-curated EQ statements used in this work are available as supplemental data of Oellrich, Walls et al. (2015). The hierarchical functional categorization of the set of Arabidopsis genes used in this work is available as supplemental data of Lloyd and Meinke (2012). The code used to produce the results of this work is available at github.com/irbraun/phenologs. Files necessary to reproduce the discussed results, datasets used to generate figures presented in this work, and other supplemental files are available at doi.org/10.5281/zenodo.3255020. This data repository also includes versions of the previously described datasets available as supplemental data of Oellrich Walls et al. (2015), and Lloyd and Meinke (2012), for the purpose of making this study reproducible without any additional external files.

## Acknowledgements

This manuscript has been released as a preprint at doi.org/10.1101/689976.

We thank Lisa Harper, Sowmya Vajjala, and Ramona Walls for helpful discussions and suggestions. We are grateful to the NSF Phenotype Ontology RCN (#DBI-0956049) for creating foundations for this work by bringing plant and computational biologists together to develop a common vocabulary and for their support to the Plant Phenotype Pilot Project participants who developed the Oellrich, Walls et al. (2015) datasets that our analyses relied upon. We thank the reviewers for their valuable guidance. Based upon their suggestions, the manuscript was improved significantly.

The authors were supported to carry out this work by an Iowa State University Presidential Interdisciplinary Research Seed Grant, the Iowa State University Plant Sciences Institute Faculty Scholars Program, and the Predictive Plant Phenomics NSF Research Traineeship (#DGE-1545453).

## Authors Contribution Statement

IRB and CJLD together contributed conception and design of the study; IRB organized the data; IRB performed the analyses; IRB wrote the manuscript; IRB and CJLD contributed to manuscript revision, read, and approved the final version.

## Footnotes

1 In relation to sentence structure, the Entity represents the subject and the Quality represents the predicate. Qualities are also elsewhere referred to as attributes, features, or characteristics of a biological structure or process.

