## Supplemental Methods and Figures for "Automated methods enable direct computation on phenotypic descriptions for novel candidate gene prediction"

#### Dataset of Phenotypic Descriptions and Curated EQ Statements

The pairwise phenotype similarity network described in Oellrich, Walls et al. (2015) was built based on a dataset of phenotype descriptions across six different model plant species (*A. thaliana*, *Z. mays* ssp. *mays*, *S. lycopersicum*, *O. sativa*, *M. truncatula*, and *G. max*). Each phenotype description was split into one or more atomized statements describing individual phenes, each of which mapped to exactly one curated EQ statement (Table 1). The EQ statements in this dataset were primarily built from terms present in PATO, PO, GO, and ChEBI, and in general follow the form

$$\text{primary } E_1 + [\text{primary } E_2] + Q + [QL] + [\text{secondary } E_1] + [\text{secondary } E_2]$$

where optional components are enclosed in square brackets. Entities are given as $E_x$ and the Quality is given as $Q$ which can be additionally described by an optional qualifier term $QL$. EQ statements were constructed within additional constraints described by Oellrich, Walls et al. (2015); for example, terms from the ChEBI ontology cannot be used to describe the first primary Entity and the Quality must be relational if any secondary entity is present.

#### Similarity Metrics

##### Jaccard Similarity

The Jaccard similarity between two sets is generally defined as the ratio of the cardinalities of the intersection and union of the sets. If $S(t)$ is defined as the set of ontology terms containing both term $t$ and all the terms that are inherited by $t$ through the ontology graph's structure, then this metric can be used to calculate the similarity of two terms $t_1$ and $t_2$ by first finding $S(t_1)$ and $S(t_2)$ and then applying the Jaccard similarity equation to these sets. Formally this equation is given as

$$J_{\text{sim}}(t_1, t_2) = \frac{|S(t_1) \cap S(t_2)|}{|S(t_1) \cup S(t_2)|} \tag{1}$$

##### EQ Statement Similarity

Similarity between EQ statements can be measured using Jaccard similarity. An EQ statement inherits all the EQ statements where the Entity and Quality are equivalent to or are a subclass of the Entity and Quality in the original statement respectively. The Jaccard similarity between two EQ statements can then be calculated between the sets of EQ statements that each inherits. We refer to this method as $\mathrm{S}_1$. EQ statements can also be treated as sets of individual ontology terms, taking the union of all terms in the statement and those they inherit. Using this approach, the Jaccard similarity between two EQ statements can then be calculated between the sets of terms representing each statement. We refer to this method as $\mathrm{S}_2$. For phenotypes for which multiple EQ statements are annotated, similarity can be calculated using $\mathrm{S}_1$ by taking the average maximal Jaccard similarity between each EQ statement in one phenotype to any EQ statement in the other.

Similarity can be calculated using $S_2$ by taking the union of the terms for all EQ statements from each phenotype and calculating the Jaccard similarity between those sets.

##### Partial Precision and Partial Recall

Partial precision and partial recall are metrics that can be used to measure the similarity between a set of target annotations, such as ontology terms assigned by curators, and a set of predicted annotations, such as those generated computationally by an algorithm (Dahdul et al., 2018). We use these metrics to evaluate the performance of methods that computationally perform semantic annotation (mapping ontology terms to input text descriptions) in order to compare the annotations made by these computational methods to the hand-curated annotations produced by Oellrich, Walls et al. (2015).

Partial precision ($PP$) is defined as the average similarity between each predicted term and the most similar corresponding target term. Corresponding terms are defined as those that were annotated to the same input text. This metric is similar to the traditional notion of binary precision because it captures the fraction of predicted information which is in fact true. Partial recall ($PR$) is defined as the average similarity between each target term and the most similar corresponding predicted term. Again, this is similar to the concept of binary recall because it captures the fraction of true information which is present in (recalled by) the predicted annotations. Formally these metrics can be given as

$$PP = \frac{1}{m} \sum_{m}^{i=1} \text{argmax}_j J_{\text{sim}}(t_i, t_j) \mathbb{1}\{y(t_i, t_j)\} \tag{2}$$

$$PR = \frac{1}{n} \sum_{n}^{j=1} \text{argmax}_i J_{\text{sim}}(t_j, t_i) \mathbb{1}\{y(t_i, t_j)\} \tag{3}$$

where $n$ is the number of target (curated) terms indexed by $j$, $m$ is the number of predicted terms indexed by $i$, $t_i$ is a particular predicted term, $t_j$ is a particular target term, and let $y(t_i, t_j)$ be true if the terms were annotated to the same input text description and false otherwise.

#### Methods for Annotating Descriptions with Ontology Terms

Semantic annotation is the process of mapping semantically-rich identifying concepts to text, or to specific words within a larger text. In the case of this work, this process involved mapping ontology terms to phenotype or phene descriptions so that these terms could be used downstream to construct EQ statements. For this task, we used NCBO Annotator, NOBLE Coder, and a Naïve Bayes bag-of-words classifier, all supported by word embedding models to account for variance in word choice. The motivation for using these tools and methods was to determine if semantic annotation could be accomplished on the dataset of phenotype and phene descriptions described in Oellrich, Walls et al. (2015) using techniques that require no manual curation.

#### NCBO Annotator and NOBLE Coder

The NCBO (National Center for Biomedical Ontology) Annotator (Musen et al., 2012) is a web service available through Bio Portal (Whetzel et al., 2011) that uses the `mgrep` concept recognition algorithm (Dai et al., 2008) to map natural language text inputs to ontology terms from a predefined set of available ontologies. The NOBLE Coder annotation tool (Tseytlin et al., 2016) is another method developed for mapping natural language inputs to ontology terms. NOBLE Coder takes user provided ontology files as reference vocabularies of synonyms and terms, and therefore can be used as a general solution for semantic annotation tasks provided only that ontologies to describe the entities of interest exist. Depending on parameter selections, this annotation tool can account for partial matches between words and ontology terms, matches with alterations in the ordering of words, and matches to multiple ontology terms that overlap in the input text. As output, both of these tools provide a set of ontology terms mapped to the text input, as well as the position within the text to which each term mapped. We used NCBO Annotator with the default set of parameters and NOBLE Coder with both precise and partial matching.

#### Naïve Bayes Classifier

A Naïve Bayes classifier was used to determine a probability $p(t|w)$ for each term $t$ and bag-of-words (un-ordered list of words) $w$ representing a given phenotype or phene description, where $p(t|w) = p(t)\prod_{i=1}^{n} p(w_i|t)$ and the bag-of-words contains $n$ individual words indexed by $i$. This value reflects the probability that $t$ should be annotated to the description given the words present in that description. The values of $p(t)$ and $p(w_i|t)$ were obtained from a fold of training data drawn from the dataset of phenotype and phene descriptions by looking at the frequency of $t$ in the curated EQ statements and co-occurrence of $t$ in an EQ statement with $w_i$ in the corresponding phenotype or phene description, respectively. Four disjoint folds of training data were used across the dataset to obtain predicted annotations for all of the phenotype and phene descriptions. In each testing data fold, the $k$ terms of an ontology $O$ with the highest values of $p(t|w)$ were output as annotations where $k$ is the average number of terms from $O$ per description in the corresponding training data multiplied by number of descriptions in the testing data. The motivation for including this method is that the Naïve Bayes classifier is capable of learning arbitrary associations between text and terms annotated to that text by curators, and unlike NCBO Annotator and NOBLE Coder, does not explicitly take into account any syntactic similarity between the text and terms.

#### Variation in Vocabulary

Word embeddings generated with Word2Vec (Mikolov et al., 2013) were incorporated to support the semantic annotation process using the above tools and methods. A model pre-trained on Wikipedia was used (Lau and Baldwin, 2016). For a particular threshold value, related words were defined as any pair of words with cosine similarity between vector representations greater than that value given the Word2Vec model. For NOBLE Coder, related words obtained through this method at a particular threshold were incorporated by inputting multiple versions of each phenotype or phene description, where additional versions were generated by iteratively substituting each word for any available related words. For the Naïve Bayes classifier, related words obtained using this method were accounted for during training when estimating the probabilities for $p(w_i|t)$. The frequency of each word $w_i$ observed in the data for a given term $t$ was increased simultaneously with the frequency of any word related to $w_i$ for that term, even though that related word was not explicitly contained within the phenotype or phene description. In this way, words were associated with terms even if those words were not explicitly present within the dataset of phenotypic descriptions.

### Constructing EQ Statements

#### Aggregating Ontology Terms

The terms annotated by each method were combined into an aggregated set of candidate ontology terms from which EQ statements were constructed. In aggregating these annotations, the union of the individual sets of annotations was taken. Terms annotated with NOBLE Coder using the precise matching parameter and NCBO Annotator were assigned a score of 1, terms annotated with NOBLE Coder using the partial matching parameters were scored based on the alignment score between the text representing the term and the portion of the description it matched, and terms annotated with the Naïve Bayes method were assigned as a score the probability $p(t|w)$ determined by the model. When aggregating terms, the maximum score for a term was retained if it was annotated by more than one method. Similarly, if two methods identified different words from the phenotype or phene description that a given term mapped to, the union of those sets of words was retained.

#### Combining Terms to Form EQ Statements

The aggregate set of ontology terms annotated to each phene or phenotype description were used to generate EQ statements. First, if no Entity terms (from PO, GO, or ChEBI) were present in the aggregated set, then the default subject term *whole plant* (PO:0000003) was added to the set of terms. If the set contained a biological process term from GO then *process quality* (PATO:0001236) was added to the set if not already present. All possible EQ statements (adhering to the format described by Oellrich, Walls et al., 2015) were then generated. From these generated EQ statements, any in which an Entity term overlapped with a Quality term were removed. (Overlapping terms in this case are defined as two terms that have an overlap in the set-of-words from the text description that was associated with those terms by the semantic annotation methods.) EQ statements were ranked by the average scores assigned during semantic annotation to ontology terms from which they were composed, then by the fraction of the words in the text description directly corresponding to an ontology term in the EQ statement. Up to the $k$ highest ranked EQ statements were retained for each text description, with $k = 4$ being the default value and used for the results described here.

### Creating Gene/Phenotype Networks

Oellrich, Walls et al. (2015) developed a network with phenotypes as nodes and similarity between them as edges for all the phenotypes in the dataset. For each type of text representations that we generated with computational methods, comparable networks were constructed. For EQ statement representations, similarity metrics described above were used to determine edge values. For vector representations generated using Doc2Vec and bag-of-words, cosine similarity was used. For the vector representations generated using set-of-words, Jaccard similarity was used. These networks are considered to be simultaneously gene and phenotype similarity networks, because each phenotype in the dataset corresponds to a specific causal gene, and a node in the network represents both that causal gene and its cognate phenotype. However, two phenotype descriptions corresponding to the same gene are retained as two separate nodes in the network, so while each node represents a unique gene/phenotype pair, a single gene may be represented within more than one node.

### Supplemental Figures

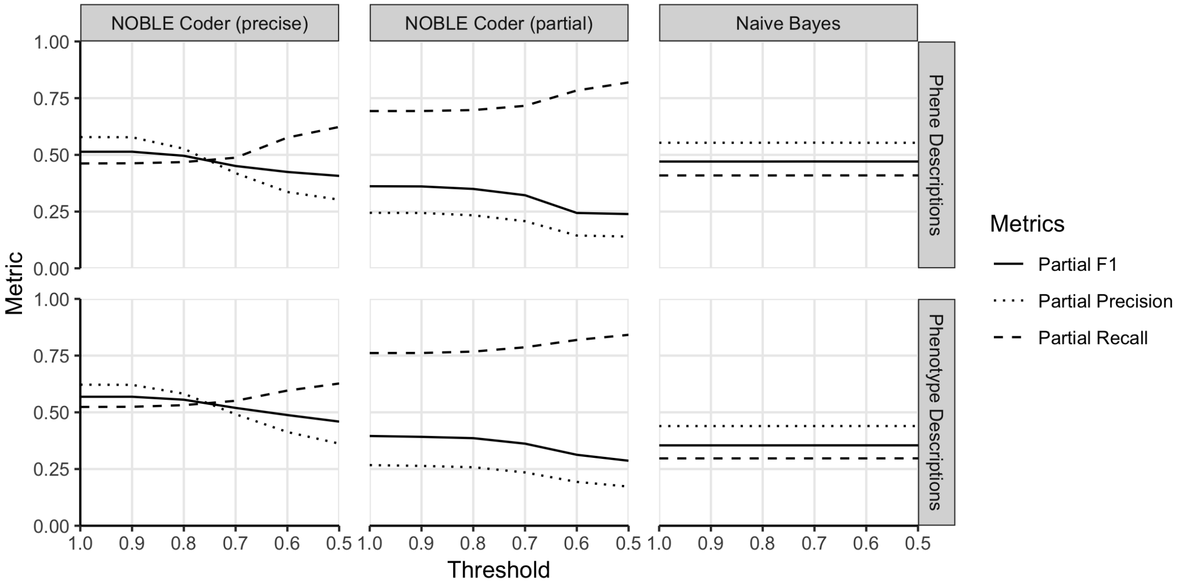


**Figure S1.** Semantic annotation metrics for methods as Word2Vec similarity threshold is decreased. The threshold indicates the similarity value at which two words are considered the same for the purposes of the semantic annotation step. The lines illustrate metrics calculated based on terms from all ontologies rather than separated by ontology.
